# Proximity Proteome Analysis Reveals Novel TREM2 Interactors in the ER-Mitochondria Interface of Human Microglia

**DOI:** 10.1101/2023.03.21.533722

**Authors:** Chulhwan Kwak, Gina M. Finan, Yu Rim Park, Anjali Garg, Oscar Harari, Ji Young Mun, Hyun-Woo Rhee, Tae-Wan Kim

**Affiliations:** Department of Chemistry, Seoul National University, Seoul 08826, Korea; The Research Institute of Basic Science, Seoul National University, Seoul 08826, Korea; School of Biological Sciences, Seoul National University, Seoul 08826, Korea; Department of Pathology and Cell Biology, Taub Institute for Research on Alzheimer’s Disease and the Aging Brain, Columbia University Irving Medical Center, New York, NY 10032, USA; Division of Biomedical science, School of Medicine, Kyungpook National University, Daegu, Republic of Korea; Neural Circuit Research Group, Korea Brain Research Institute, 41062 Daegu, Korea; Department of Psychiatry, Washington University School of Medicine, St. Louis, MO 63108, USA

## Abstract

Triggering receptor expressed on myeloid cells 2 (TREM2) plays a central role in microglial biology and the pathogenesis of Alzheimer’s disease (AD). Besides DNAX-activating protein 12 (DAP12), a communal adaptor for TREM2 and many other receptors, other cellular interactors of TREM2 remain largely elusive. We employed a ‘proximity labeling’ approach using a biotin ligase, TurboID, for mapping protein–protein interactions in live mammalian cells. We discovered novel TREM2-proximal proteins with diverse functions, including those localized to the Mitochondria-ER contact sites (MERCs), a dynamic subcellular ‘hub’ implicated in a number of crucial cell physiology such as lipid metabolism. TREM2 deficiency alters the thickness (inter-organelle distance) of MERCs, a structural parameter of metabolic state, in microglia derived from human induced pluripotent stem cells. Our TurboID-based TREM2 interactome study suggest novel roles for TREM2 in the structural plasticity of the MERCs, raising the possibility that dysregulation of MERC-related TREM2 functions contribute to AD pathobiology.

## Introduction

Alzheimer’s disease (AD) is the leading cause of dementia in the elderly. In addition to plaques, tangles, and widespread neuronal death, there are profound inflammatory changes in the AD brain, which are increasingly presenting new therapeutic targets (Fu et al., 2019; Wyss-Coray and Mucke, 2002). Microglia act as the brain’s immune cells, representing roughly 10% of the cells in the brain, and are becoming the subject of intense investigation in the field of neurodegenerative disease research (Leng and Edison, 2021; Hansen et al., 2018; Sarlus et al., 2017). In AD, microglia are linked closely to all pathological cascades including amyloid, tau and neuroinflammatory changes (McQuade and Blurton-Jones, 2019). Recent genome-wide association studies (GWAS) revealed that a number of AD risk genes are highly (and often exclusively) expressed in microglia, including the gene encoding triggering receptor expressed on myeloid cells-2 (TREM2) (Colonna, 2023; Villegas-Llerena et al., 2016; Griciuc and Tanzi 2021; Jorfi et al., 2023). Notably, a rare R47H hypomorphic mutation in the TREM2 increases the risk of AD several fold (R. Guerreiro et al., 2013; Jonsson et al., 2013). Gene network studies reveal that TREM2 and its adapter protein, DNAX-activation protein 12 (DAP12; also known as TYROBP) strongly associate with AD (Zhang et al., 2013; 2016; Matarin et al., 2015). Homozygous variants in TREM2 or its binding partner DAP12 are also shown to cause Nasu-Hakola disease (NHD) with early-onset dementia (Paloneva et al., 2002; Klunemann et al., 2005). Despite the strong clinical phenotype of human NHD, TREM2-deficient and DAP12-deficient mice do not exert strong NHD-like phenotypes (Zhou et al., 2023).

TREM2 is a transmembrane immune signaling receptor, which serves as a receptor for β-amyloid (Zhao et al., 2018; Lessard et al., 2018; Zhong et al., 2018) and various lipoproteins including high density lipoprotein containing apolipoprotein E (Song et al., 2017; Atagi et al., 2015; Bailey et al., 2015). TREM2 promotes the survival and proliferation of microglia and mediates the clearance of apoptotic neurons via phagocytosis without inducing apparent neuroinflammation (Takahashi et al., 2005). TREM2 is essential for retaining a fully mature disease-associated microglia (DAM) profile and for supporting the microglial response to Aβ-plaque-induced pathology (Keren-Shaul et al., 2017; Krasemann et al., 2017). TREM2-dependent response requires the induction of the phosphorylation of tyrosine residues within the ITAM motifs of DAP12, by Src-family kinases and downstream signaling cascades including AKT and mTOR (Kober et al., 2017; Jay et al., 2017). Despite extensive investigation, we currently do not have a complete mechanistic understanding of microglial TREM2 signaling, due in part to the lack of a comprehensive knowledge of TREM2-bearing molecular complex(es) in microglial cells. Most of the focus has been on the interaction of TREM2 with DAP12, which serves as a communal platform for downstream signaling. In addition to TREM2, DAP12 has been shown to serve as a common hub that interacts with numerous receptors (Lanier, 2009; Haure-Mirande 2022). Given the complexity and high disease relevance of TREM2-associated biological processes, it is conceivable that additional molecular components may interact with TREM2 and contribute substantially to the regulation of relevant microglial functions.

We set out to identify novel components of TREM2-harboring protein complex(es), not yet implicated in TREM2 signaling, using an unbiased *in situ* proteomics approach. To isolate protein complexes of interest and analyzed the components present in the complex, conventional procedures involve detergent lysis, subcellular fractionation, and affinity pull-down followed by mass spectrometry, which often produces false positives or fails to discover weak or unstable interactions (Wilson and Nairn 2018; Mehta and Trinkle-Mulcahy 2018; Bludau and Aebersold 2020). We used TurboID-based “proximity labeling (PL)” technique to investigate the protein-protein interactions in living microglial cells (Lam et al., 2015; Han and Ting, 2018; Lee et al., 2019; Qin et al., 2021; Choi and Rhee, 2022). TurboID, an engineered biotin ligase, uses ATP to convert biotin into biotin-AMP, a reactive intermediate that covalently labels biotin to the proximal proteins (Doerr, 2018; Brandon et al., 2018). Recently, we improved the TurboID-based PL further and performed ‘super resolution PL’, which involves the highly selective capture of biotinylated peptides, allowing the detection the TurboID-mediated biotinylation event at the single amino acid level (Lee et al., 2023).

Using the human microglial cells stably expressing TurboID-fused to the cytoplasmic region of TREM2, we performed super resolution proximity labeling experiments followed by mass spectrometry analysis, and successfully identified and validated 47 putative TREM2 interacting proteins. Interestingly, among these proteins, 21 proteins are found to localize to the subcellular compartment known as the mitochondria-ER contacts (MERCs) (Vance 1990; 2014; Lee et al., 2018). Biochemically isolated MERCs are also referred to as mitochondrial associated membranes (MAMs; Vance 2014). MERCs have been implicated in many functional roles associated with TREM2, including lipid transport, mitochondrial energy metabolism, Ca2+ handling and immune sensing (Rieusset, 2018; Tubbs and Rieusset, 2017), and have also been suggested to be a pathological hub for several neurodegenerative diseases, including AD (Area-Gomez et al., 2018; Bose and Beal, 2016; Paillusson et al., 2016; Johri and Chandra, 2021; Area-Gomez and Schon, 2017). Given the physical and/or functional association of TREM2 to MERC-related cellular activities, we investigated the effects of TREM2 deficiency on the cell biological changes of MERCs in the human induced pluripotent stem cells (iPSC)-derived microglia (iMGL) (Abud et al., 2017; McQuade et al., 2018). TREM2 deficiency leads to the ultrastructural alteration of MERCs, further supporting a functional association of TREM2 to MERCs. Our finding also raises the possibility that TREM2-mediated microglial functions relevant to AD may be attributed to the aberrant regulation of MERCs in microglia.

## Results

### Proximity labeling activity of TREM2-TurboID in human microglial cell line

For enzyme-based proximity labeling (PL), we selected TurboID, one of the engineered biotin ligases that is suitable for protein-protein interaction mapping due to the high catalytic activity as compared to other PL enzymes (Branon 2018). We generated mammalian expression plasmids encoding human TREM2 fused to a C-terminal V5 tag and TurboID (TREM2-V5-TurboID) (Fig. 1A). TREM2-V5-TurboID was transfected into the human microglial cell line, IM-HM (Mazzeo et al., 2022). Western blotting with antibodies against V5-tag or TREM2 revealed that TREM2-V5-TurboID is expressed with the predicted size (Fig. 1B). Confocal fluorescent microscopy showed that TREM2-V5-TurboID localizes mainly to intracellular compartments as previously reported (Figures 1C; Kleinberger 2014; Park 2015; Yin 2016; Zhao 2017). We next performed confocal microscopy analysis to detect subcellular distribution of TREM2-V5-TurboID-mediated biotinylation using streptavidin-647 (Fig. 1C). TREM2-V5-TurboID immunoreactivity and biotin-labeled signals appear to co-localize strongly, suggesting that the biotin labeling occurs at subcellular compartments in close proximity to TREM2-V5-TurboID. TurboID conjugated to nuclear export signal and V5 tag (V5-TurboID-NES) was used as a control, and the biotin signals from V5-TurboID-NES vs TREM2-V5-TurboID occurred in spatially distinct subcellular sites (fig. S1A). We next biochemically validated PL activity of TREM2-V5-TurboID in IM-HM cells. Upon incubation with exogenous biotin, a wide range of biotinylated proteins were detected in TREM2-V5-TurboID-expressing cells (fig. S1B). The biotin-labeled proteins were then analyzed by Western blotting using streptavidin-HRP. Distinct patterns of biotinylated proteins were observed in the samples from the V5-TurboID-NES-expressing IM-HM cells (fig. S1C).

**Fig. 1.**
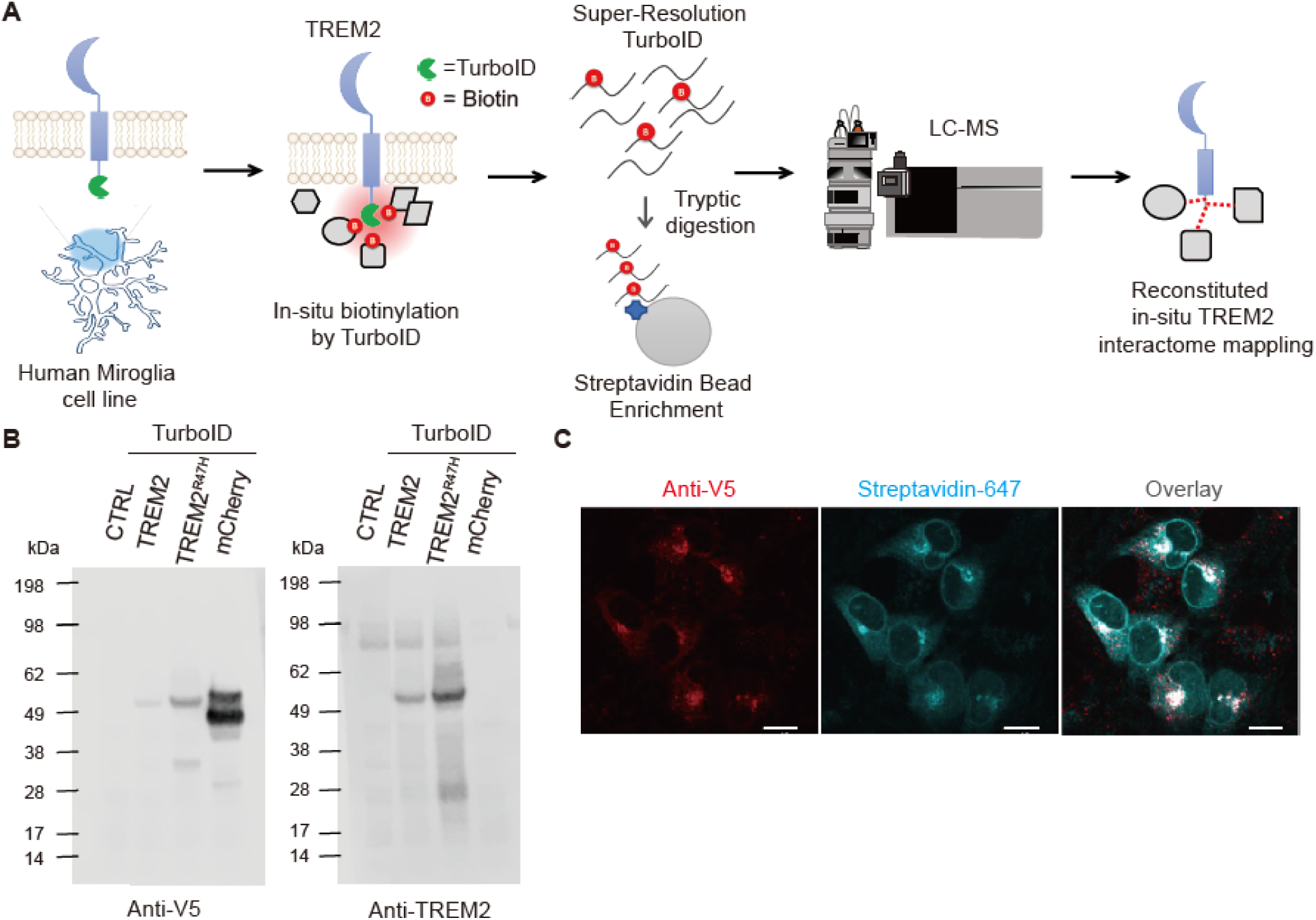
TurboID-based proximity labeling of TREM2 in microglial cells. (**A**) Schematic workflow of proximity labeling, super-resolution TurboID, mass spectrometry and in situ TREM2 interactome mapping. (**B**) Western blot analysis of the samples from the IM-HM microglial cells transfected with the expression constructs: human wild-type or R47H version of TREM2 with C-terminal V5 tag and TruboID and mCherry-V5. (**C**) Confocal microscopy analysis of IM-HM cells transfected with TREM2-V5-TurboID using anti-V5 antibodies or with Streptavidin-647 for the detection of biotin-labeled protein.

### Proximal proteomics of TREM2 in living microglial cells

We established stable IM-HM transfectants expressing TREM2-V5-TurboID and performed ‘super resolution PL’ (Fig. 1A; Lee at al. 2023). The samples were subjected to streptavidin pull-down to enrich biotin-conjugated proteins. We then performed on-bead trypsin digestion, allowing direct mass spectrometry analysis of the peptide harboring biotin-conjugated lysine (K+226) (Lee et al., 2023). This approach removes the bulk of unlabeled peptides, therefore drastically improving detection sensitivity and reducing false positive results. After verifying the biotin labeling, biotin-labeled peptides were analyzed by mass spectrometry as previously described (Lee et al., 2019; 2023; Kwak et al., 2020). By statistically assessing the relative enrichment of the biotinylated peptides from TREM2-V5-TurboID relative to the control (V5-TurboID-NES), we identified 47 proteins as components of a TREM2 interactome (Fig. 2A).

**Fig. 2.**
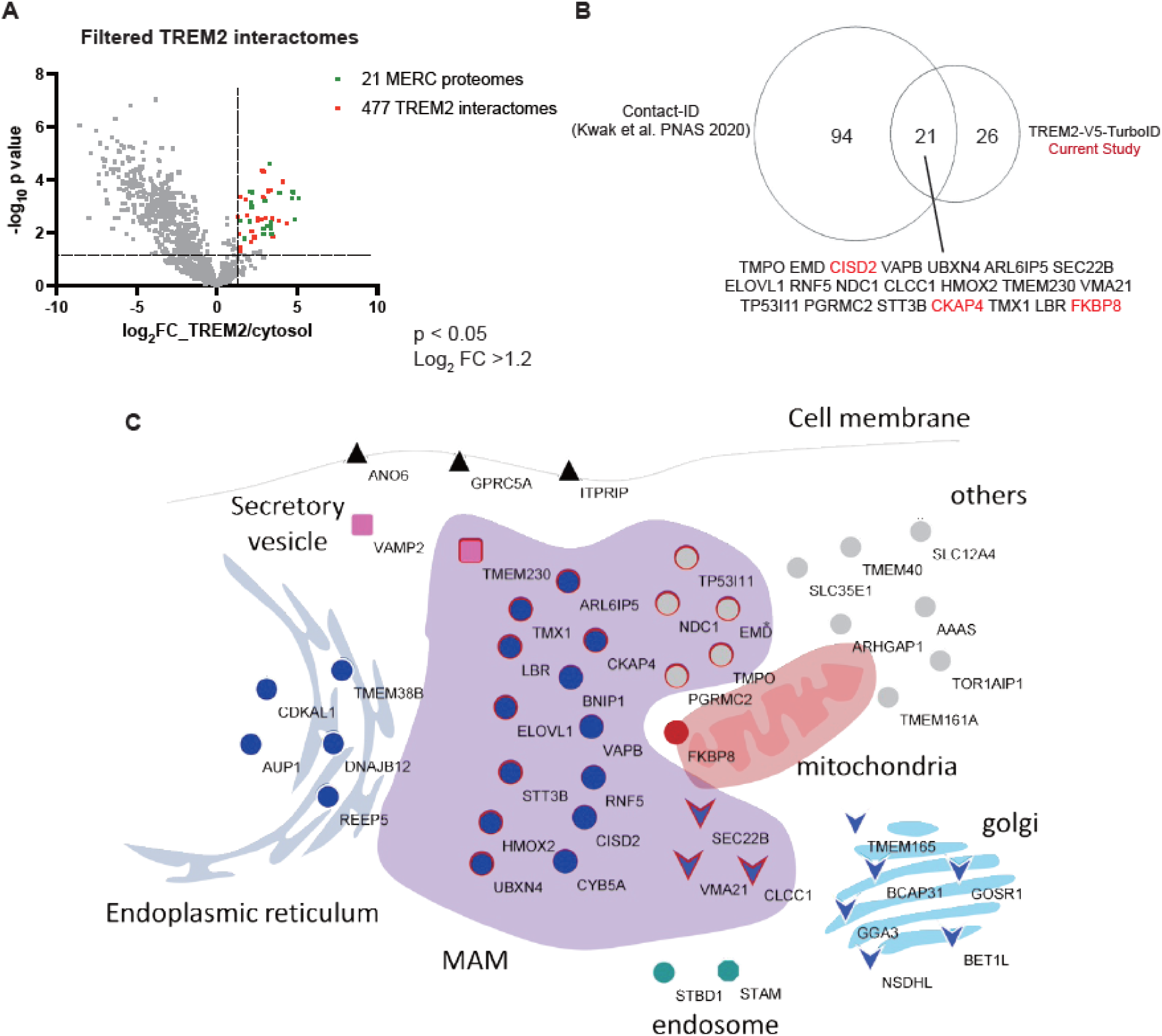
Clustering of TREM2 interactome. (**A**) Volcano plots showing statistically significant enrichment of biotinylated proteins (group-TREM2-TurboID) over the cytosolic biotinylated proteins by NES-TurboID (TREM2 interactome). Among the TREM2 interactome, MERC components are also indicated. Enrichment of the Spot-TurboID detected biotinylated proteins by TREM2-V5-TurboID. x-axis: Log2 value of fold change (FC) of mass signal intensities. Cut off value is 1.2; y-axis: -Log10 value of p-value, cut off value is 1.2. (**B**) Segregation of TREM2 interactome vs. previously known MERC components shown by Contact-ID (Kwak et al., 2020). (**C**) Subcellular distribution of TREM2 interactome proteins across the MERC (MAM), ER, mitochondria, secretory vesicles, Golgi and cell surface.

### Analysis and validation of TREM2 interactome

Quantification of biotinylated peptides derived from the TREM2 proximal proteins was performed as previously described (Lee et al., 2019). Our TREM2 interactome analysis revealed a total of 47 proteins (Fig. 2A). To our surprise, among the 47 identified proteins, we found that 21 TREM2-proximal proteins are known components of MERCs (Fig. 2, A and B). These MERC components were identified in a previous study via ER-OMM contact-dependent localization studies between ER and the outer mitochondrial membrane (OMM) (Kwak et al., 2020). Quantitative proteomics allowed us to determine the MS signal intensity of MERC components vs. non-MERC components (fig. S2). Our assessment of the TREM2-TurboID interactome reveals novel TREM2 interactors in the MERCs of a human microglial cell line. We summarized the TREM2 interactome and their corresponding subcellular sites (Fig. 2C). To validate the physical proximity of TREM2-V5-TurboID and identified proteins, we performed a proximity-dependent biotinylation assay using the expression plasmid encoding each “hit” protein (fig. S2A). An expression construct encoding TREM2-V5-TurboID was co-transfected with each expression “hit” construct in 293 cells and subjected to in situ biotinylation assay for validation. This assay successfully validated the interaction between TREM2-V5-TurboID and the well-known TREM2 interactor, DAP12, as well newly discovered interactors such as CISD2 and CKAP4 (fig. S1B). PL-mediated biotin incorporation is shown to be quantitative (Han and Ting, 2018; Choi and Rhee, 2022). DAP12 exhibited the highest levels of biotin incorporation as compared to other identified components of TREM2 interactome (fig. S1C). Our 47 confirmed TREM proximal proteins were found to be positive. We need to add the description for old fig S2 (new fig. S3)!

### TREM2 localization to the MERCs

We next investigated the subcellular localizations of TREM2-proximal proteins. To precisely map the MERCs, we used ‘Contact-ID’, in which two inactive fragments of BioID (a PL enzyme), are reconstituted via protein-protein interaction across two distal membranes and become catalytically active (Fig. 3A; Kwak et al., 2020). The Contact-ID system consists of the N-G78 (B1) fragment of BioID conjugated to the cytosol-facing N terminus of SEC61B (B1-SEC61B) at the ER membrane and the G79-C (B2) fragments of BioID conjugated to the C terminus of TOM20 (TOM20-B2) at the OMM. This approach allows for precise mapping of the MERC contact sites without overlapping distribution of virtually any MERC components with either the ER or OMM. We transfected TREM2-V5 without TurboID and examine the subcellular distribution of TREM2 vs. Contact-ID-dependent signals (Fig. 3B). Confocal microscopy analysis showed that the subcellular localization pattern of TREM2 overlaps substantially to that of the Contact-ID signal. Biochemical data showed that TREM2 is indeed proximal to the MERC components (Fig. 3C). Please add additional description on F3C here. These data suggest that TREM2 localizes to the MERCs and biotinylation of TREM2 is mediated by the PL activity of TREM2-TurboID within the MERCs.

**Fig. 3.**
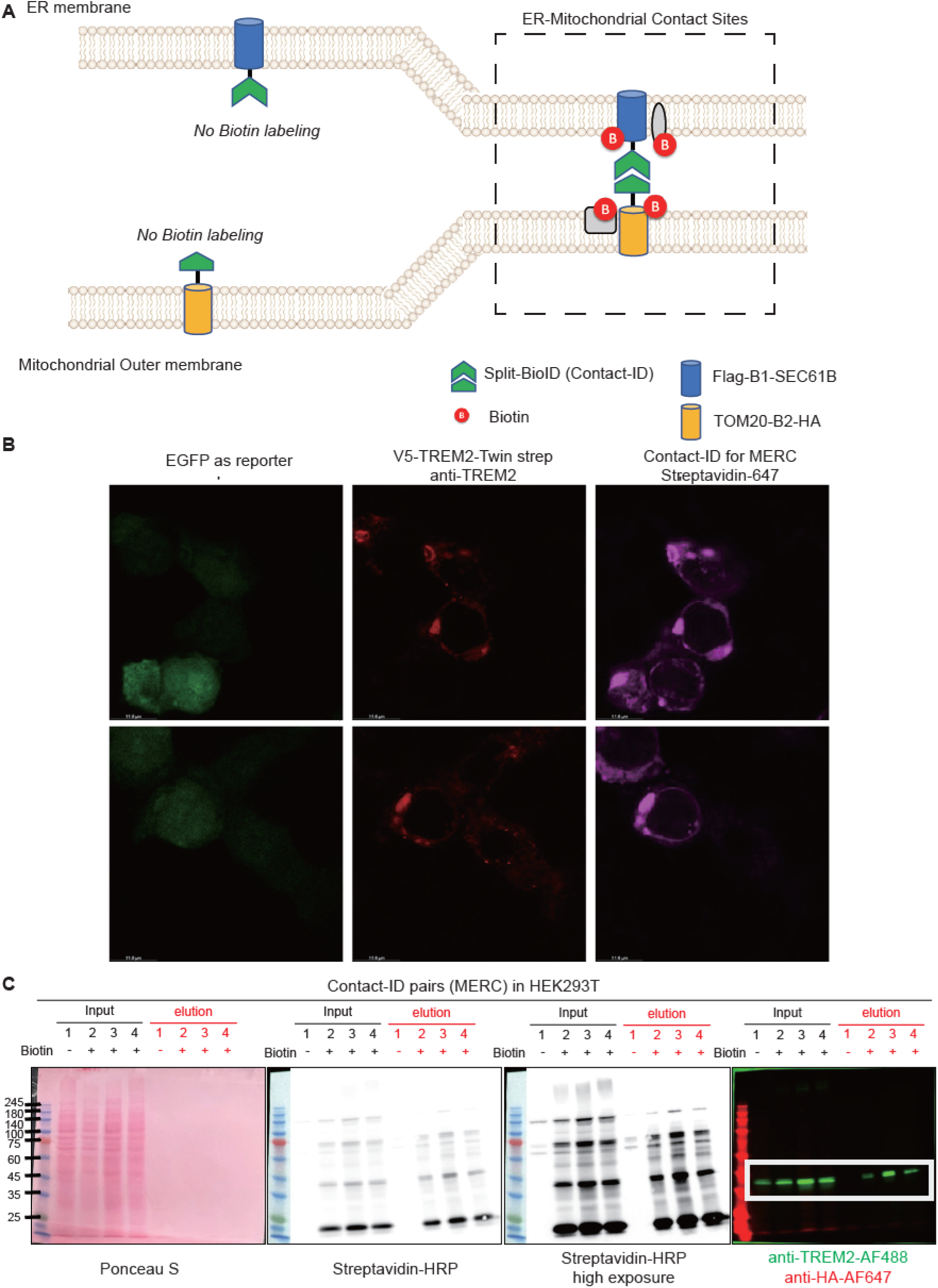
Demonstration of MERC localization of TREM2 using the contact-dependent PL labeling. (**A**) Schematic diagram of Contact-ID for MERC mapping using split BioID. (B) Confocal microscopy imaging of MERC biotinylation by Contact-ID in HEK293T cells. TREM2 was visualized by anti-V5 antibody. Biotinylated proteins were visualized by AF647-conjugated streptavidin. (**C**) Streptavidin-horseradish peroxidase (SA-HRP) Western blot of biotinylated proteins.

### TREM2 deficiency increases the thickness of MERCs in iMGL

Prevalent localization of the TREM2 interactome in the MERCs suggests that TREM2 may be required for the proper structural and/or functional regulation of MERCs in human microglia. Recently, human induced pluripotent stem cell (iPSC)-derived microglial models (iMGL) for AD became available (Abud et al., 2017; McQuade et al., 2018). We therefore investigated the cell biological impact of TREM2 deficiency in human iMGL -a more physiological model system as compared to a microglial cell line. We performed electron microscopy analysis to test if TREM2 deficiency affects the ultrastructure of MERCs in iMGL (Fig. 4). Consistent with the previous mouse *in vivo* studies (Nugent et al., 2020), there is a marked increase of lipid droplets (LDs) in TREM2-deficient iMGL as compared to wild-type iMGL (fig. S4). We next performed morphological characterization of MERCs in wild-type vs. TREM2 deficient iMGL (Fig. 4A). The established key structural parameter of MERCs includes the width of the gap (10-30 nm distance) between cytosolic sites of ER membrane and OMM (MERC thickness; Csordás et al., 2006; Giacomello and Pellegrini, 2016). MERC thickness is a regulated parameter strongly associated with metabolic state of the cells. Cell function and survival relies on the maintenance of a proper spacing between the ER and mitochondria (Csordás et al., 2006; Giacomello and Pellegrini, 2016). We found that overall MERC thickness is greatly increased in TREM2 knockout iMGL as compared to wild-type (Fig 4, B and C). To ensure that the observed increases of the MERC thickness in the TREM2-deficient iMGL is specific to cell biological alterations owing to TREM2, we next tested whether re-introduction of TREM2 into the TREM2-deficient iMGL could restore the observed MERC alterations associated with TREM2 deficiency (Fig. 5). Lentiviral delivery of TREM2 in the TREM2 deficient iMGL leads to a decrease in the MERC thickness as compared to iMGLs infected with control lentiviral particles (Fig. 5). In addition, the MERC thickness increased occurrence of LDs, a prominent phenotype of TREM2-deficient iMGL, was also rescued by the introduction of wild-type human TREM2 (fig. S5). These data suggest that the increased MERC thickness in the TREM2-deficient cells is a TREM2-specific cell biological change.

**Fig. 4.**
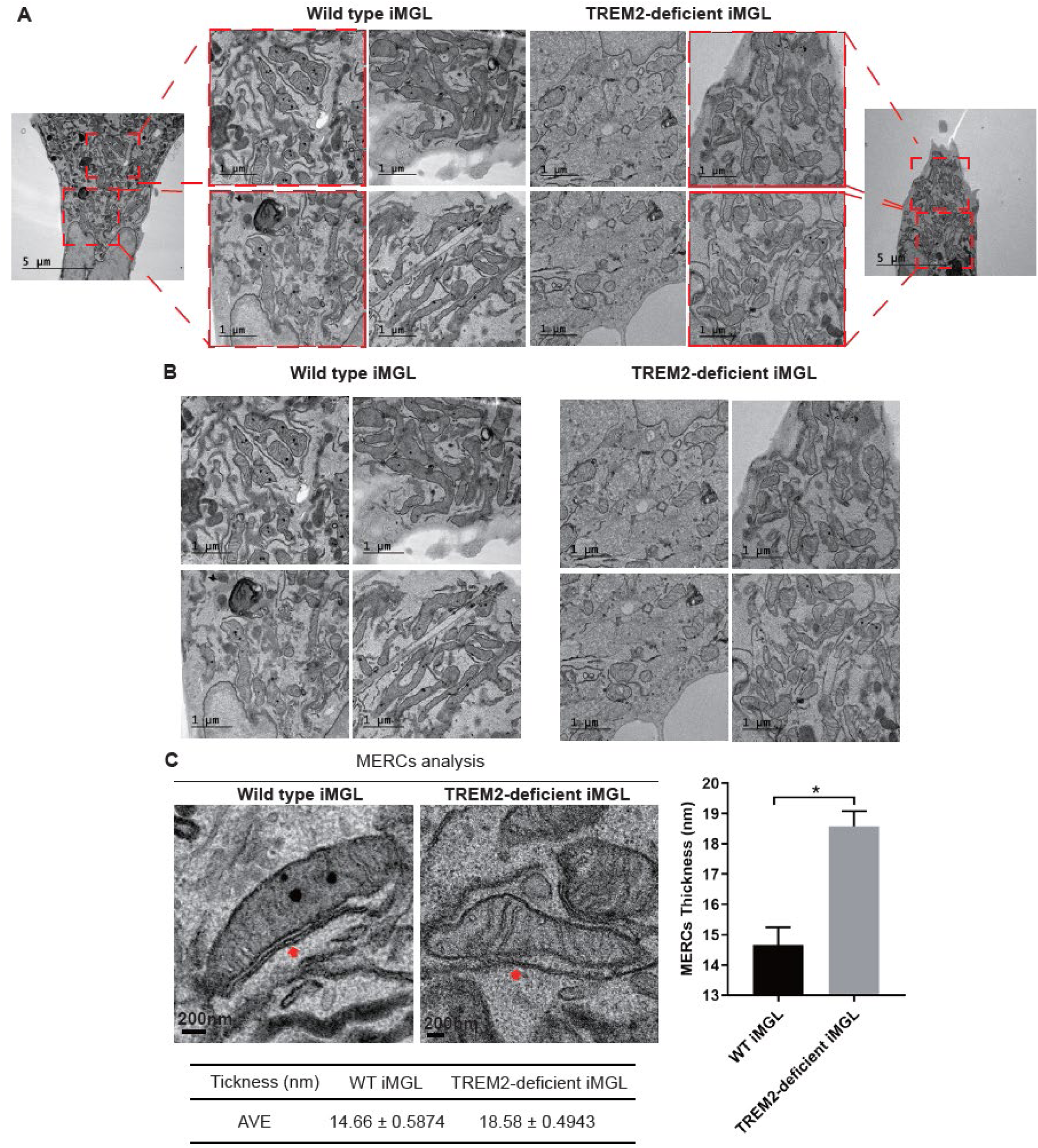
Ultrastructural Analysis of MERCs in wild type iMGL and TREM2 deficient iMGL. (**A**) Transmission Electron microscopy (TEM) images of wild-type and TREM2 deficient iMGL cells. Red boxes are enlarged images of dotted area. Each ultrastructure of intracellular organelles including mitochondria and ER is intact without abnormality. scale bar: 1 um. (**B**) The thickness and length of MERC were analyzed in each group. Red arrows represent MERC. The thickness of MERCs with less than 30 nm between the mitochondria and ER was measured in 10 regions of MERCs. (**C**) The averaged thickness was significantly increased in TREM2 deficient iMGL, and the thickness (nm) was each 14.66 ± 0.5824 and 18.58 ± 0.4943 in wild-type and TREM2 deficient cells. N=50 MERCs. Data are represented as mean ± SEM and the statistical significance was p <0.001.

**Fig. 5.**
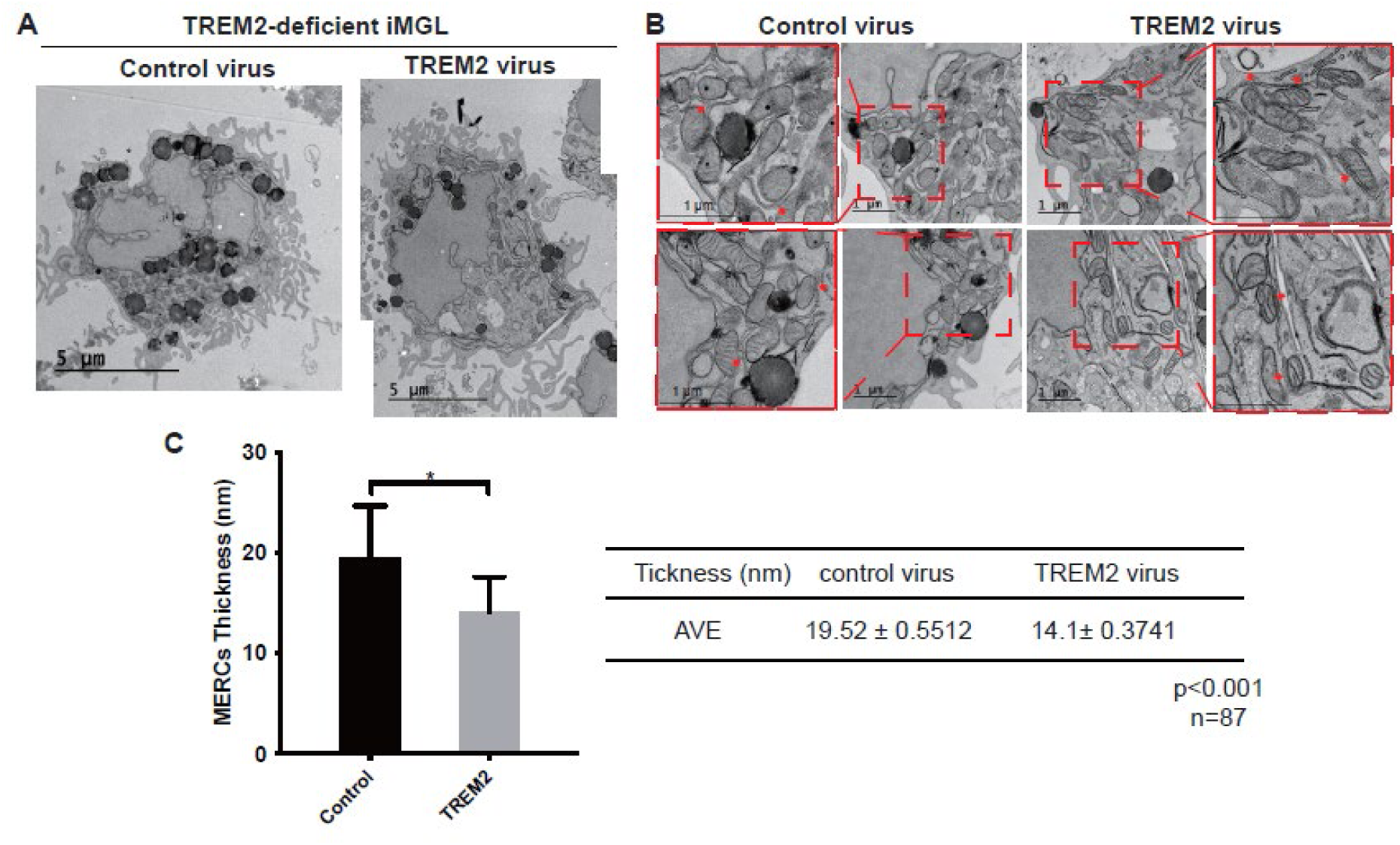
The thickness of MERC in TREM2 deficient iMGL and the iMGL infected with TREM2 lentivirus. (**A**) Transmission Electron microscopy (TEM) images of TREM2 deficient iMGL cells with control lentivirus (left), and TREM2 lentivirus (right). An increased number of LDs in TREM2-deficient iMGL was prominent, in contrast, LD was decreased in the cell infected with TREM2 lentivirus. (**B**) Black squares are enlarged images from dotted rectangles showing MERC. Arrows showed MERC. (**C**) Increased thickness of MERC in TREM2 deficient iMGL was rescued by TREM2 encoding vector. Each averaged thickness (nm) of MERC was 19.52 ± 0.5512 and 14.1 ± 0.3741 in TREM2 deficient iMGL and the cell infected with TREM2 encoding vector. N=87 MERCs. Data are represented as mean ± SEM and the statistical significance was p <0.001.

### Effects of R47H TREM2 on the MERC thickness

The loss-of-function variants of TREM2, including R47H, are associated with an increased risk of AD (Guerreiro et al., 2013; Jonsson et al., 2013). To test whether the altered MERC thickness is associated with hypomorphic TREM2 R47H variant, we next analyzed the MERC thickness of the iMGL derived from human iPSC carrying the R47H TREM2 variant (Figure 6A). R47H TREM2 loss of function

**Fig. 6.**
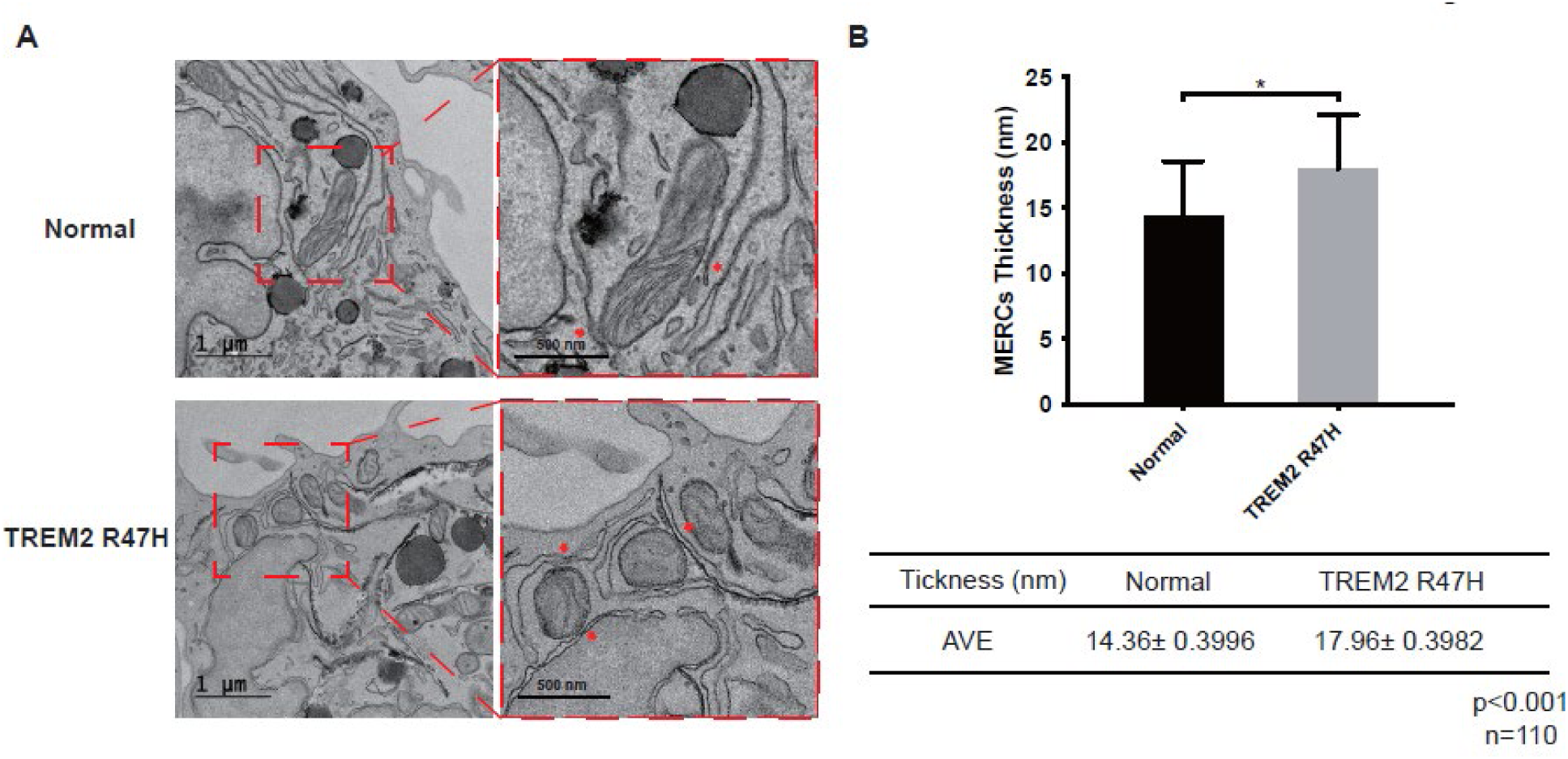
Increased thickness of MERC in TREM2 R47mutated iMGL. (A) Transmission Electron microscopy (TEM) images of iMGL cells from wild type and patient with R47H mutation. Enlarged images from dotted boxes (Right) showed MERCs. (B) The thickness of MERCs was analyzed in isogenic wild type and TREM2 R47H mutated iMGL. Each averaged thickness (nm) of MERC was 14.36 ± 0.3996 and 17.96 ± 0.3982 in isogenic wild type and TREM2 R47H mutated iMGL. N=110. Data are represented as mean ± SEM and the statistical significance was p <0.001.

We determined the distance of ER-to-mitochondria of iMGL carrying TREM2 R47H mutant, as compared to isogenic wild-type control. The MERC thickness was significantly elevated in the TREM2 R47H iMGL relative to control (Fig. 6B). The MERC thickness phenotype was less profound than that observed in the TREM2 full knockout iMGL, probable due to the partial loss-of-function characteristics of TREM2 R47H (Cheng-Hathaway et al., 2018; Sudom et al., 2018). The occurrence of LDs was also increased in the iMGLs carrying R47H TREM2 as compared to isogenic wild-type (fig. S5).

### Enrichment of TREM2 interactome genes across microglia transcriptional states

To assess potential involvement of the TREM2 interactome with AD, we next leveraged the single-cell RNA-sequencing (scRNA-seq) data from human AD samples. Specifically, we analyzed whether the transcriptomic profiles of the genes encoding the MERC TREM2 interactome associate with specific activation state(s) among transcriptionally distinct sub-populations of human microglia, including homeostatic, activated, IFN and IL1B (Fig. 7). The scRNA-seq microglia data was collected from human donors (parietal cortex from 67 donors from the Knight ADRC and DIAN brain banks; N=16,849) (Brase et al., 2023). Curated marker genes were employed to label the cell types based on the overall expression profile of the nuclei as previously described (Da Masquita et al., 2021). Detailed clinical data and genetic characterization of the collected microglia are described in Table 1. Here, we used four different microglia subtypes based on their transcriptomic profiles and corresponding functional/activation status: mic-resting (n=7001), mic-activated (n=3653), mic-reduced (n=2078) and mic-inflammatory (n=1668) (Fig. 7, A and B). The average expression of the genes encoding the TREM2 interactome in MERC (MERC genes) into each microglia transcriptional states by Control vs. TREM2 reduced (R47H, R62H, H157Y variants) genetic status was done (Fig. 7C). A MERC genes score was also calculated for microglia cell by running Seurat’s AddModuleScore. The enrichment of TREM2 interactome MAM genes into Control and TREM2 reduced microglia cell was also evaluated (Fig. 7D). Further, the significant pairwise differences into each Control and TREM2 reduced transcriptional states was calculated using the linear mixed model. moduleScore ∼ Cluster + (1|Subject) (Fig. 7E).

**Fig. 7.**
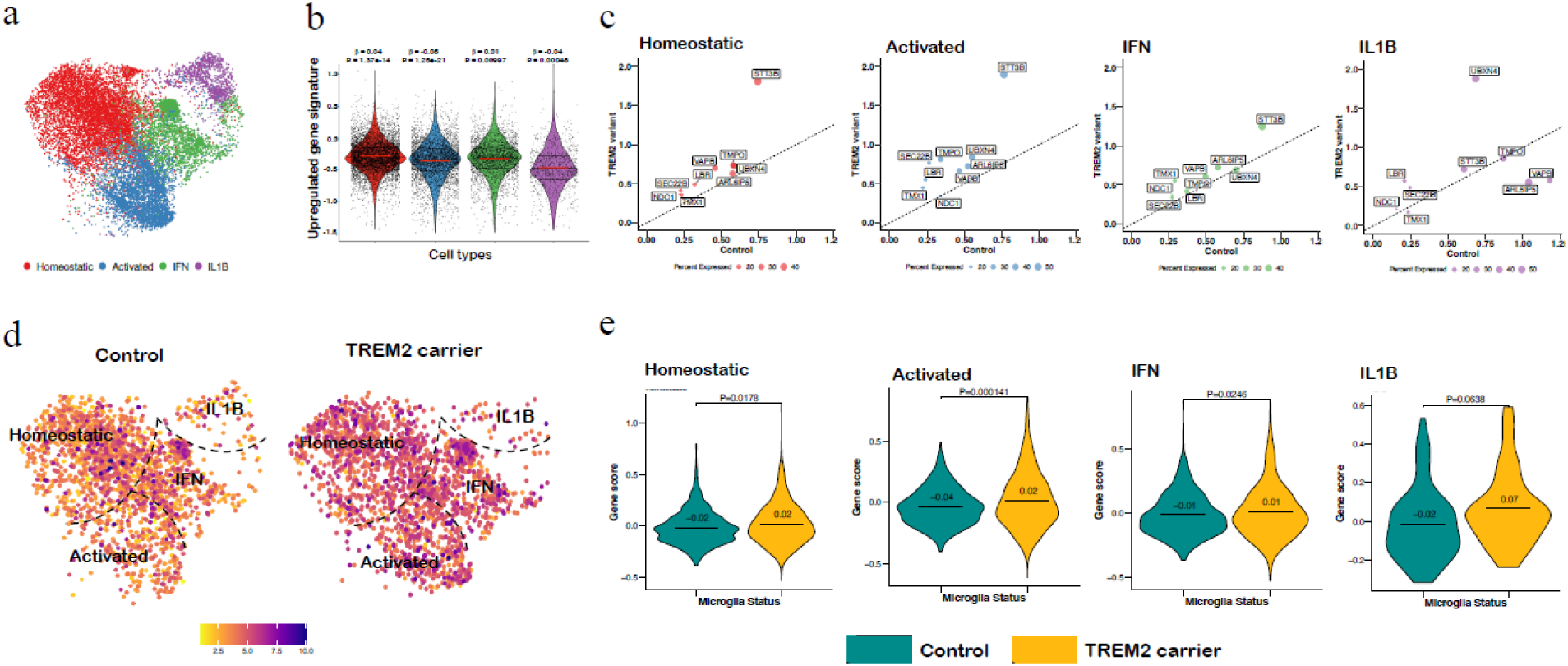
Expression of MERC proteins in human microglia derived from control and TREM-carrier of R47H, R62H, and H157Y mutation. (**A**) UMAP plot showing the microglia nuclei from the Homeostatic (overexpression of TMEM119, p= 3.34 x 10^-5^), Activated (overexpression of CD9, p = 2.93E-93), IFN (overexpression of IFIT3, p= 3.24 x 10^-58^), and IL1B (overexpression of IL1B, p= 8.98 x 10_-200_) clusters. (**B**) Expression of MERC gene in microglia transcriptional state. Red line shows the mean expression value. (C) Differential gene expression of each MERC genes (scale expression >0.1) in the four microglia transcriptional states. (**D**) UMAP representation of the Control (N = 2251; left) and TREM2-carrier (N = 2136; right) cells. The color code represents the expression profile of the MERC genes (depicted in panel c) in each microglia transcriptional state. (**E**) Comparison of expression profile of MERC gene in Control and AD carrier in TREM2-carrier risk variants.

## Discussion

Protein interaction networks underlie the biological processes and pathological progression of human diseases (Rees et al., 2015; Bludau and Aebersold, 2020). Given the critical role for TREM2 in microglial biology as well as the pathophysiology of AD, a comprehensive understanding of the protein-protein interaction networks of TREM2 should provide new insights on TREM2 functions in microglia. We utilized an TurboID-catalyzed PL approach combined with mass spectrometry to determine the TREM interactome in live microglial cells. The PL approach overcomes some inherent limitations of the traditional methods associated with affinity pull-down or co-immunoprecipitation, which often involve a post-lysis manipulation of the starting material (e.g. detergent extraction, subcellular fractionation). PL can detect native protein-protein interactions in live cells, and is thereby highly suitable for mapping target protein interactome with a greater specificity. Among a number of PL enzymes, TurboID was chosen because of its catalytic efficiency, simplicity and the use non-toxic biotin substrate (Branon et al., 2019). Here we identified novel proteins proximal to TREM2-TurboID, including those localized predominantly to MERCs, unveiling the connection between TREM2 and MERCs.

MERC is a dynamic and plastic structure that can undergo remodeling in response to metabolic change and stress (Rowland 2012; Csordás et al., 2006). The thickness of MERC, the width of the cleft that separates the ER and OMM, is considered a regulated structural parameter that changes depending on the metabolic state of the cells (Giacomello and Pellegrini, 2016). Changes in the width of the cleft in MERC influence the lipid transfer process across two membranes (Giacomello and Pellegrini, 2016). We have demonstrated that deficiency or hypomorphic R47H mutation of TREM2 lead to increased thickness of MERCs in iMGs (Fig. 4 and 6). In addition to our novel observation of the MERC alterations, our results show the increased occurrence of LDs in TREM2 deficient iMGL (fig. S4). This is consistent with the previous report demonstrating the increased accumulation of cholesterol ester (e.g. CE20:4 and CE22:6), the major cholesterol metabolites stored in LDs, in both mouse brain and cultured human myeloid cells (Nugent et al., 2020). Since MERC has been shown to play an important role in the biogenesis of LD (Liu et al., 2015; Freyre et al., 2019; Choudhary et al., 2021; Guyard et al., 2022), it is conceivable that altered MERC in the TREM2 deficient iMGL may be attributed to the aberrant accumulation of LDs in iMGL. When cell conditions promote active expansion of LDs, the synthesis and trafficking of new phospholipids are regulated in the area between mitochondria and ER (Olzmann et al., 2019). According to a report by Area-Gomez et al., the ACAT protein relating LD generation was more abundant in the mitochondria-associated ER membranes (MAMs) fraction and had approximately 3 times higher enzymatic activity compared to the ER and mitochondrial fractions (Area-Gomez et al., 2012). Collectively, virtually all known TREM2 functions including metabolic regulation of lipids and Ca2+, as well as inflammatory activation, are linked intimately to those of MERC, substantiating the idea that TREM2 is an important factor for proper MERC functions.

Aberrant MERCs have been reported in cellular models of AD. Gene deletion and FAD mutations of familial AD-associated presenilin genes (PSEN1 and PSEN2) greatly increases the MERC thickness (50-200 nm) in fibroblasts (Area-Gomez et al., 2012). Presenilin 2 has been shown to regulate Ca2+ crosstalk across the MERCs (Zampese et al., 2011). PS1 has also been shown to interact with TREM2 (Zhao et al., 2017). In addition to AD, MERCs have been implicated in other neurodegenerative diseases, such as Parkinson’s disease (PD) (Area-Gomez et al., 2018; Bose and Beal, 2016; Paillusson et al., 2016; Johri and Chandra, 2021; Area-Gomez and Schon, 2017). Occurrence of the microglial TREM2 interactome in the MERCs is particularly relevant to a known MERC functions involving immune activation. MERC is shown to serve as a signaling hub for NLRP3 inflammasome activation (Horng et al., 2014; Zhou et al., 2020). TREM2 is a direct regulator of the NLRP3 inflammasome and the AD-linked R47H TREM2 variant is not capable of activating the inflammasome (Cosker et al., 2021; Li et al., 2021; Jiang et al., 2022; Wang et al., 2022). These observations further substantiate the idea of functional association of TREM2 with MERC.

In our previous study, we developed Contact-ID, a split-pair BioID system used to identify the protein components of MERCs (Kwak et al., 2020). Since all known MERC markers show overlapping subcellular distribution in either the ER or OMM, the Contact-ID approach allows precise mapping of ER-OMM contact-dependent localization (Fig. 3), indicting co-localization of TREM2 with identified MERC components in the TREM2 interactome. In addition to the components of MERC, our PL interactome study revealed newly discovery TREM2-interacting proteins with distinct subcellular localizations (Fig. 2C), suggesting potential mediators or modulators of cellular functions of TREM2 in microglial cells. For instance, anoctamin 6 encoded by ANO6 is an essential component for Ca^2+^-dependent exposure of phosphatidylserine (PS) on the cell surface (Yang et al., 2012; Kunzelmann et al., 2014). PS exposure is considered to be one of the “eat-me” signals for the engulfment by immune cells (Lemke et al., 2019; Scott-Hewitt et al., 2020) and ANO6 has been shown to increase microglial phagocytosis of neurons (Batti et al., 2016; Zhang et al., 2020). Interestingly, ANO6 deficiency reduces activation of the microglial NLRP3 inflammasome in the brain of AD mouse model (Cui et al., 2023), suggesting the functional link of ANO6 with TREM2 in the regulation of NLRP3 inflammasome (Li et al., 2021; Jiang et al., 2022). CDK5 regulatory subunit Associated protein 1-Like 1 (CDKAL1) has been implicated in the pathogenesis of type2 diabetes (Ghosh et al., 2022) and a family-based GWAS meta-analysis revealed genetic association of two SNPs in the CDKAL1 gene (Herold et al., 2016). ITPRIP gene (The gene encoding inositol 1,4,5-trisphosphate receptor interacting protein) has also been suggested as an AD risk gene (Simino et al., 2017). AUP1 and REEP5 are known to play a key role in LD biogenesis (Robichaud et al., 2021; Chen et al., 2022), consistent with the functional role for TREM2 in the LD biogenesis. Further studies are needed for the characterization of the components of the TREM2 interactome in microglial cells, including other non-MERC proteins. Future studies would also need to elaborate if the observed TREM2-associated MERCs occurs in vivo especially in the context of mouse models. Collectively, our current study provides new insights on TREM2 functions and serves as a foundation for further research aimed at deciphering new molecular players in microglial TREM2 functions.

## Materials and Methods

### Culture of iMGL

The wild type iMGL, TREM2 deficient iMGL, and iMGL derived from human iPSC carrying R47H TREM2 variant (Fujifilm Cellular dynamics iCell microglia, AHN, TREM2 R47H, #11969) were cultured in iCell microglia media kit (Fujifilm cellular dynamics, USA, #R1204) as recommended by a manufacturer. Cells were maintained on 0.07 % Poly ethyleneimine solution coated plate in suspension. For re-introduction of TREM2 into the TREM2-deficient iMGL, TREM2-deficient iMGL was infected with two different lentiviral particles (Control lentiviral particles (CH404_empty_GFP reporter_pDual) and TREM2 delivery lentiviral particles (CH400_ss-V5-TREM2-twin strep tag_GFP reporter_pDual)).

### Transmission electron microscopy (TEM) analysis

Cells were pre-fixed with 2% paraformaldehyde/2.5% glutaraldehyde and post-fixed in 2% Osmium tetroxide (OsO4; EMS, USA). After block staining with 1.5% potassium ferrocyanide (sigma, USA), 1% thiocarbohydrazide (TCH; TCI, Japan), 2% OsO4, and 2% uranyl acetate (EMS, USA), dehydration with a graded ethanol series (20%, 50%, 70%, 90%, and 100%) and infiltration with an epoxy resin (30, 50, 70, and 100%) were done. Then, samples were embedded with an EMBed-812 embedding kit (EMS, USA) following the manufacturer’s instructions. Cells were sectioned at 60 nm using an ultramicrotome (UC7; Leica Microsystems, Germany) and sections were mounted on copper slot grids with a formva film. The sections were imaged by the Tecnai G2 microscope with a US1000X-P camera 200 (Thermo Fisher Scientific, USA) at an accelerating voltage of 120 kV.

### Analysis of MERCs in iMGL

Ultrastructural analysis of MERCs in iMGL was performed as previously described (Giacomello et al., 2016; Raturi et al., 2016; Bartok et al., 2019) The thickness of MERCs with less than 30 nm between the mitochondria and ER was measured in 10 regions of MERCs. MERCs thickness and length in each group were determined in nanometers and averaged. Data were analyzed using a Image J software. Statistical analysis was performed using GraphPad Prism 8 software (GraphPad Software Inc., LaJolla, CA). The statistical significance was p <0.001.

### Analysis of lipid droplets in iMGL

To identify the number of lipid droplets (LDs), cells were stained with Lipi-red (Dojindo, Japan; 1 umol/L) and Hoechst solution (Sigma, USA; 1 umol/L). Live cells were imaged under a confocal light microscopy (Ti-RCP, Nikon, Japan). The ultrastructure of LDs was observed under a TEM and the number of LDs was measured by Image J software.

## Author contributions

H.-W.R and T.-W.K. conceived the study and wrote the manuscript. C.K., G.M.F., Y.R.P., J.Y.M., H.-W.R and T.-W.K. designed the experiments. C.K., G.M.F., Y.R.P. performed experiments and analyzed and interpreted the data. A.G. and O.H. analyzed and interpreted bioinformatics data.

## Data and materials availability

All data need for evaluating the main conclusions of this work are included in the paper and/or Supplementary Materials.

